# Remote-Controlled Wireless Bioelectronics for Fluoxetine Therapy to Promote Wound Healing in a Porcine Model

**DOI:** 10.1101/2024.12.18.629272

**Authors:** Houpu Li, Narges Asefifeyzabadi, Kaelan Schorger, Prabhat Baniya, Maryam Tebyani, Alexie Barbee, Hsin-ya Yang, Wan Shen Hee, Anthony Gallegos, Kan Zhu, Cynthia Recendez, George Luka, Sujung Kim, Koushik Devarajan, Tiffany Nguyen, Sydnie Figuerres, Celeste Franco, Elham Aslankoohi, Min Zhao, Roslyn Rivkah Isseroff, Mircea Teodorescu, Marco Rolandi

## Abstract

Wound healing presents a significant challenge in biomedical science, requiring precise therapeutic delivery and real-time monitoring. Bioelectronic systems offer a promising solution but remain largely unexplored for wound care, particularly in large animal models that reflect human healing dynamics. This study introduces a remote controlled wireless bioelectronic platform equipped with an iontophoretic pump to deliver fluoxetine, a selective serotonin reuptake inhibitor that promotes wound repair. In vitro and ex-vivo testing validated efficient on demand fluoxetine delivery. In vivo experiments in a porcine wound model demonstrated clear therapeutic efficacy over 3-day and 7-day periods. The system enhanced healing outcomes, increasing re-epithelialization by 37% (H&E staining), reducing the M1/M2 macrophage ratio by 33%, and stimulating neuronal growth at the wound site. This bioelectronic platform delivers fluoxetine in a controlled, remotely-controlled manner while allowing for wound direct wound imaging that can be used to monitor wound healing progress. Additionally, it allows precise dose and temporal delivery of treatment to enhance the outcome of future large animal wound healing studies.

## Introduction

Wound healing is a multifaceted biological process crucial for restoring tissue integrity and function following injury. [1] This intricate process involves a coordinated cascade of cellular and molecular events, including inflammation, tissue formation, and remodeling. [2] Effective management of both chronic and acute wounds remains a significant biomedical challenge, often complicated by factors such as infection, impaired vascularization, and underlying medical conditions. [3] The economic burden of wound care reaches billions of dollars annually. [4] Traditional methods, such as passive bandages and manually administered drugs, are often ineffective in addressing the dynamic and evolving needs of the wound healing process. [5] These limitations underscore the necessity for innovative strategies that provide effective therapeutic delivery and real-time insights into wound progression. [6]

Emerging technologies integrating controlled drug delivery with real-time wound monitoring can improve wound care. [6-7] Such approaches hold promise for delivering personalized and adaptive treatments, optimizing healing outcomes, and reducing healthcare costs. [8] Among potential therapeutic agents, fluoxetine, a selective serotonin reuptake inhibitor (SSRI), has shown remarkable potential beyond its established role in treating depression. [9] Recent studies indicate that fluoxetine can promote tissue regeneration, enhance neuronal growth, and reduce inflammation in wound models. [10] Controlled and targeted delivery of fluoxetine in wounds is challenging due to inherent limitations in drug delivery systems. [11] Current methods often lack the precision, programmability, and adaptability required to address the dynamic nature of wound environments. [12]

Bioelectronic address these challenges with electronic components that interact directly with biological tissue and enable precise control over therapeutic delivery. [13] Among bioelectronic systems, ion pump actuators have emerged as a promising technology for targeted drug delivery. [14] These actuators use voltage to drive ionic or molecular species, providing a programmable and precisely targeted method for delivering therapeutics. [14b, 15] These characteristics make bioelectronic delivery an ideal candidate for personalized wound care. [16] Here, we present a remote-controlled wireless bioelectronic platform capable of delivering fluoxetine (Flx) in a large animal porcine wound model (Fig. 1 A). This platform is designed to allow direct imaging of the wound with an optional camera for continuous monitoring of the healing platform (Fig. 1B). Deployment of this platform on a porcine model indicates that Flx treatment increases wound re-epithelization and reduces wound inflammation as measured by ration between pro-inflammatory (M1) and pro-reparative (M2) macrophages.

**Figure 1.**
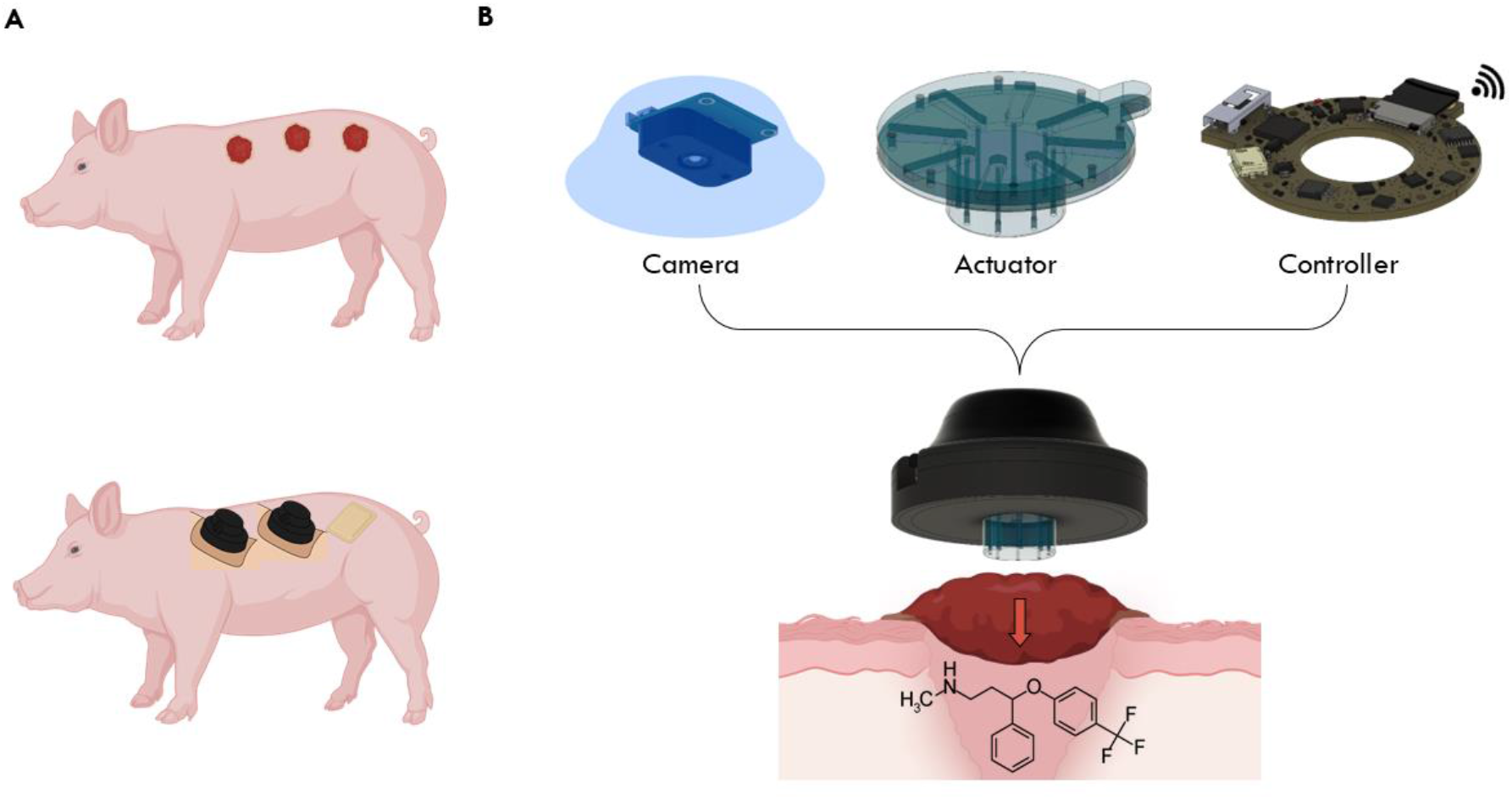
Design of the bioelectronic system for wound healing. (A) Application of the bioelectronic device on a porcine wound model to accelerate healing. (B) schematic illustration of the device components, including a bioelectronic ion pump actuator and integrated electronics for wireless monitoring of delivery rates and wound progression in a large animal model. The ion pump actuator employs voltage-driven mechanisms to enable programmable, targeted, and controlled delivery of fluoxetine cations directly to the wound site.

### Design and Fabrication of the Bioelectronic Device

The bioelectronic device integrates a multi-channel ion pump actuator and a wireless imaging module into a compact and wearable format, designed to enhance both functionality and usability (Figure 1B). The ion pump employs voltage-driven actuation to deliver fluoxetine ions directly to the wound area with precision and programmability. This targeted delivery approach ensures optimal therapeutic effects while minimizing off-target distribution.

The actuator features a polydimethylsiloxane (PDMS)-based structure that incorporates reservoirs and microchannels for controlled drug delivery. These reservoirs are coupled with hydrogel ion exchange membranes, which serve as the interface between the device and the wound. The use of PDMS ensures biocompatibility and structural stability, while the hydrogels enable efficient ion exchange and therapeutic delivery. This innovative design also maintains optical clarity, facilitating unobstructed imaging of the wound site.

A key aspect of the device is its modular design, which supports wireless control and real-time data transmission. An integrated printed circuit board (PCB) enables seamless communication between the device components, ensuring efficient operation and data management (Figure 1B). The wireless imaging module captures high-resolution images of the wound, providing continuous monitoring capabilities. This dual functionality, precise therapeutic delivery, and real-time imaging position the device as a significant advancement in wearable bioelectronics for wound care

The design and fabrication of the bioelectronic device follows a bottom-up approach to ensure precision and integration of key functionalities (Fig. 2). The core of the device is the bioelectronic ion pump actuator, which operates based on voltage-driven mechanisms to deliver fluoxetine ions to the wound site in a controlled manner (Fig. 2A). This actuator utilizes a polydimethylsiloxane (PDMS) framework, which houses reservoirs and microchannels for the delivery process (Fig. 2B).

**Figure 2.**
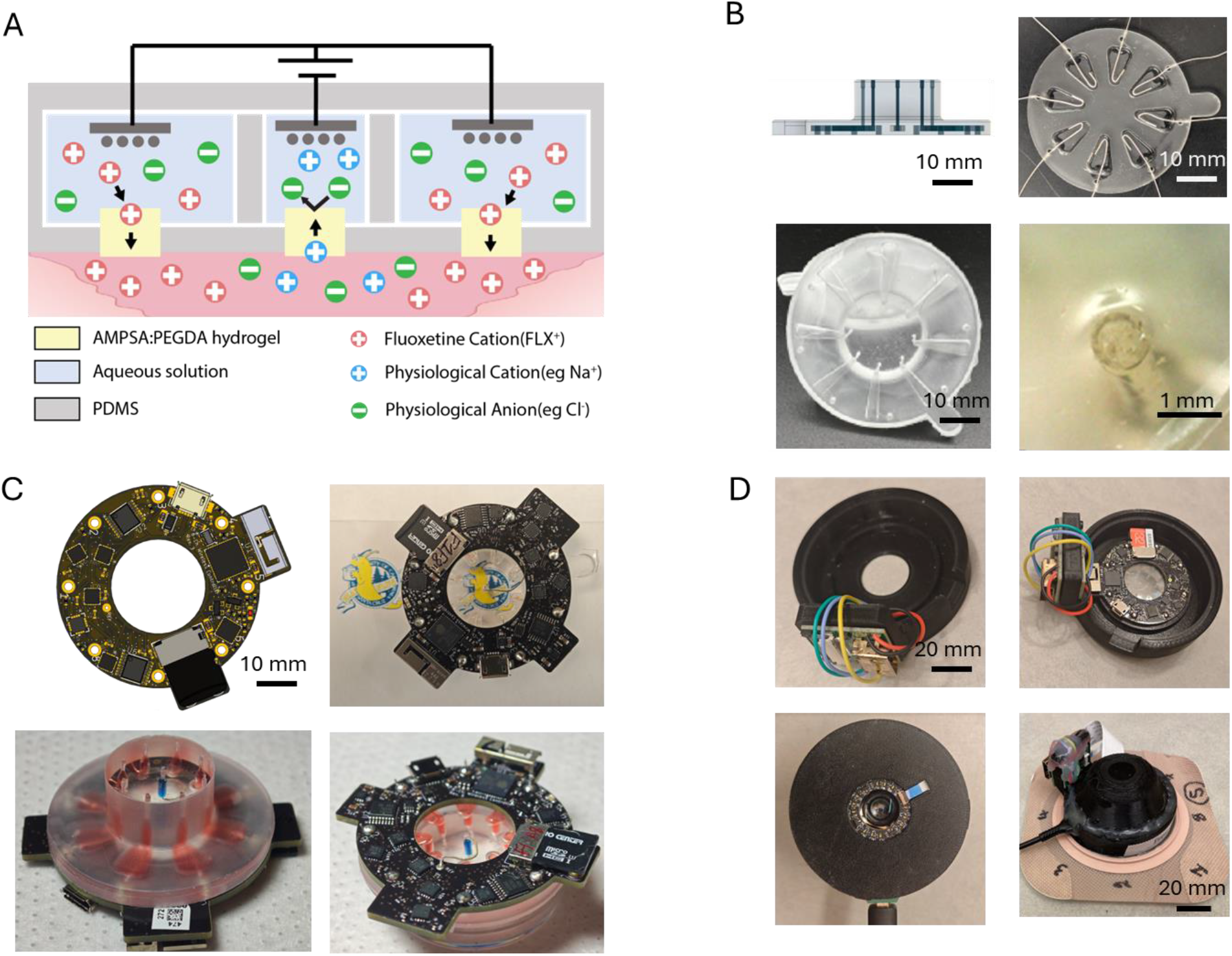
Design and bottom-up fabrication of the bioelectronic device. (A) Schematic illustration of the working principle of the bioelectronic ion pump actuator, showcasing voltage-driven delivery of Flx ions. (B) Detailed depiction of the PDMS-based actuator structure, including reservoirs and microchannels for drug delivery. Hydrogel ion exchange membranes are cured within glass capillaries embedded in the PDMS, providing a biocompatible interface with the wound for efficient drug delivery. (C) Integration of the PDMS actuator with a printed circuit board (PCB) that facilitates wireless communication and voltage control. An optical image of the device with red food dye highlights the reservoir design and optical clarity for imaging. (D) Final assembly of the bioelectronic device, enclosed for secure placement on the wound, incorporating a camera module for real-time imaging and monitoring of wound healing progress.

To enhance functionality, the actuator incorporates hydrogel ion exchange membranes cured within glass capillaries embedded in the PDMS structure (Fig. 2B). These membranes serve as the interface with the wound, enabling efficient and localized drug delivery. The design ensures compatibility with the wound environment while maintaining biocompatibility and stability.

The PDMS structure is integrated with a PCB, which facilitates wireless communication[17] and provides the driving voltage for the actuator channels (Fig. 2C). This modular integration allows for real-time control and monitoring of the therapeutic delivery process. The optical clarity of the PDMS ensures that imaging from the top is unobstructed, enhancing the diagnostic capabilities of the device. The design is validated by adding red food dye to the reservoirs, highlighting the visibility and functionality of the PDMS channels during operation.

The assembled bioelectronic device is enclosed in a protective casing, which houses additional components, such as the camera module [18] for imaging (Fig. 2D). This enclosure enables secure placement of the device on the wound while ensuring durability and ease of use.

Figure 3 demonstrates the in vitro performance of the bioelectronic device, including drug delivery simulations, high-performance liquid chromatography (HPLC) analysis, and imaging validation, which collectively establish its functionality and reliability. The delivery efficiency of the ion pump was evaluated through a series of in vitro experiments. Fluoxetine was delivered into a solution under various voltage conditions, showcasing precise control over dosage and delivery rates across eight independent channels (Fig. 3B). These experiments highlight the device’s ability to maintain consistent and programmable therapeutic outputs.

**Figure 3.**
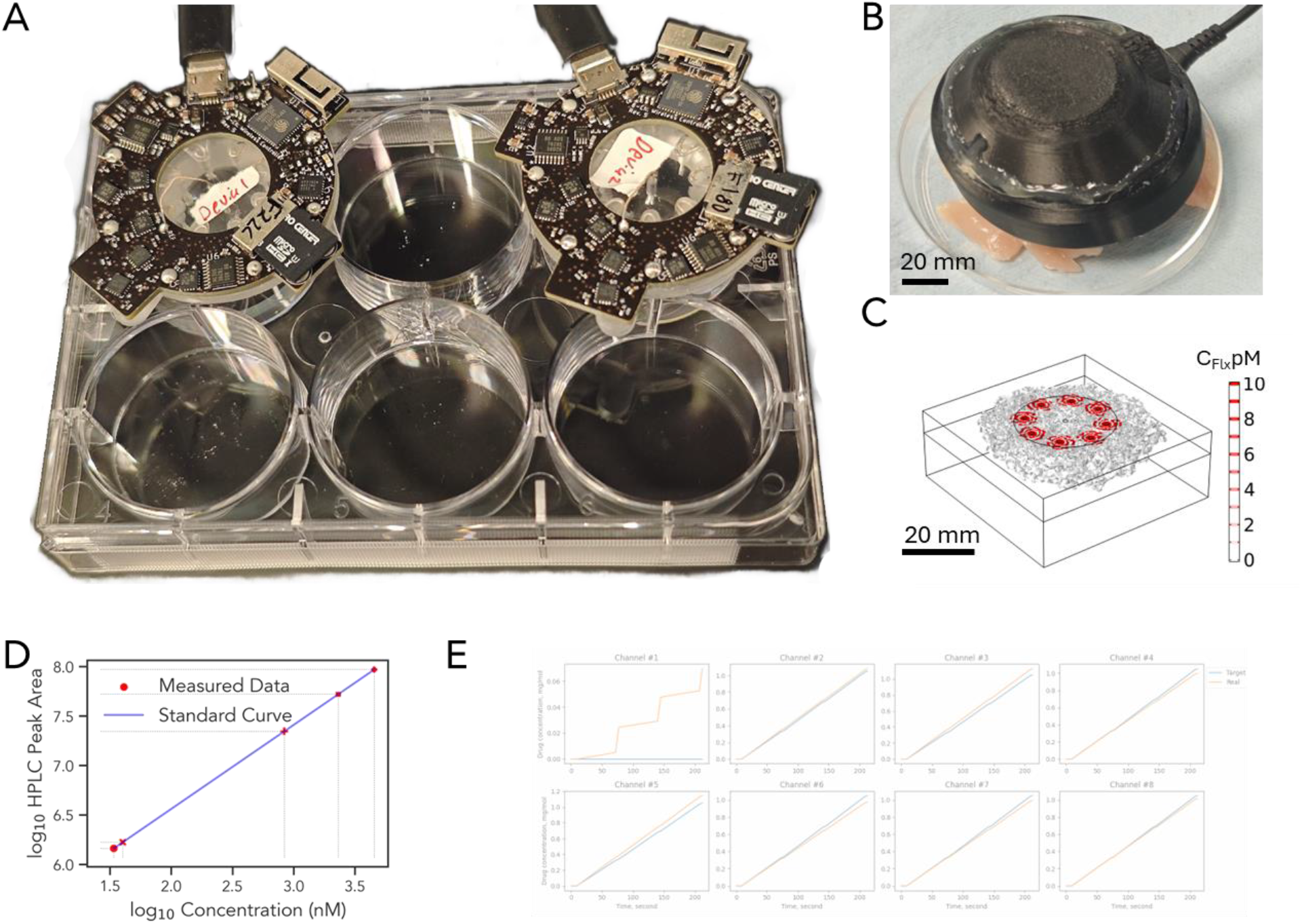
In vitro validation of the bioelectronic device. (A) The bioelectronic device tested for in vitro drug delivery performance into a solution under controlled conditions. (D) Validation of device performance using chicken breast tissue as a wound-tissue mimic to evaluate its interface with biological substrates. (C) Simulation of fluoxetine delivery from eight independent ion pump channels, demonstrating precise control over dosage and delivery rates. (D) High-performance liquid chromatography (HPLC) analysis confirming the consistency and accuracy of fluoxetine delivery, correlating with programmed current inputs. (E) Functional performance of the device actuator, showcasing effective operation in delivering of fluoxetine.

HPLC analysis further confirmed the accuracy of fluoxetine delivery. The analysis demonstrated a strong correlation between the delivered drug concentrations and the programmed current inputs, ensuring reliability in drug administration (Fig. 3C).

Validation on biological substrates, specifically chicken breast tissue as a wound-tissue mimic, provided additional evidence of the performance of the device. These tests verified the actuator’s ability to interface effectively with biological substrates while maintaining precise fluoxetine delivery (Fig. 3D-E).

Imaging tests validated the camera module’s functionality. High-resolution images of the wound area were captured through the optically clear PDMS enclosure, confirming the capacity of the device to monitor therapeutic processes without visual obstructions.

These results collectively underscore the capability of the bioelectronic device to deliver therapeutics with precision and monitor wound conditions in real-time, paving the way for advanced wound care solutions.

The in vivo experimental setup demonstrated the integration of the bioelectronic device with large animal models, specifically porcine wound models, as outlined in Figure 4. The experimental schedule included both 3-day and 7-day timelines to assess short-term and extended performance. The study evaluated the device’s performance in promoting wound healing over both short (3-day) and extended (7-day) periods (Fig. 4A).

**Figure 4.**
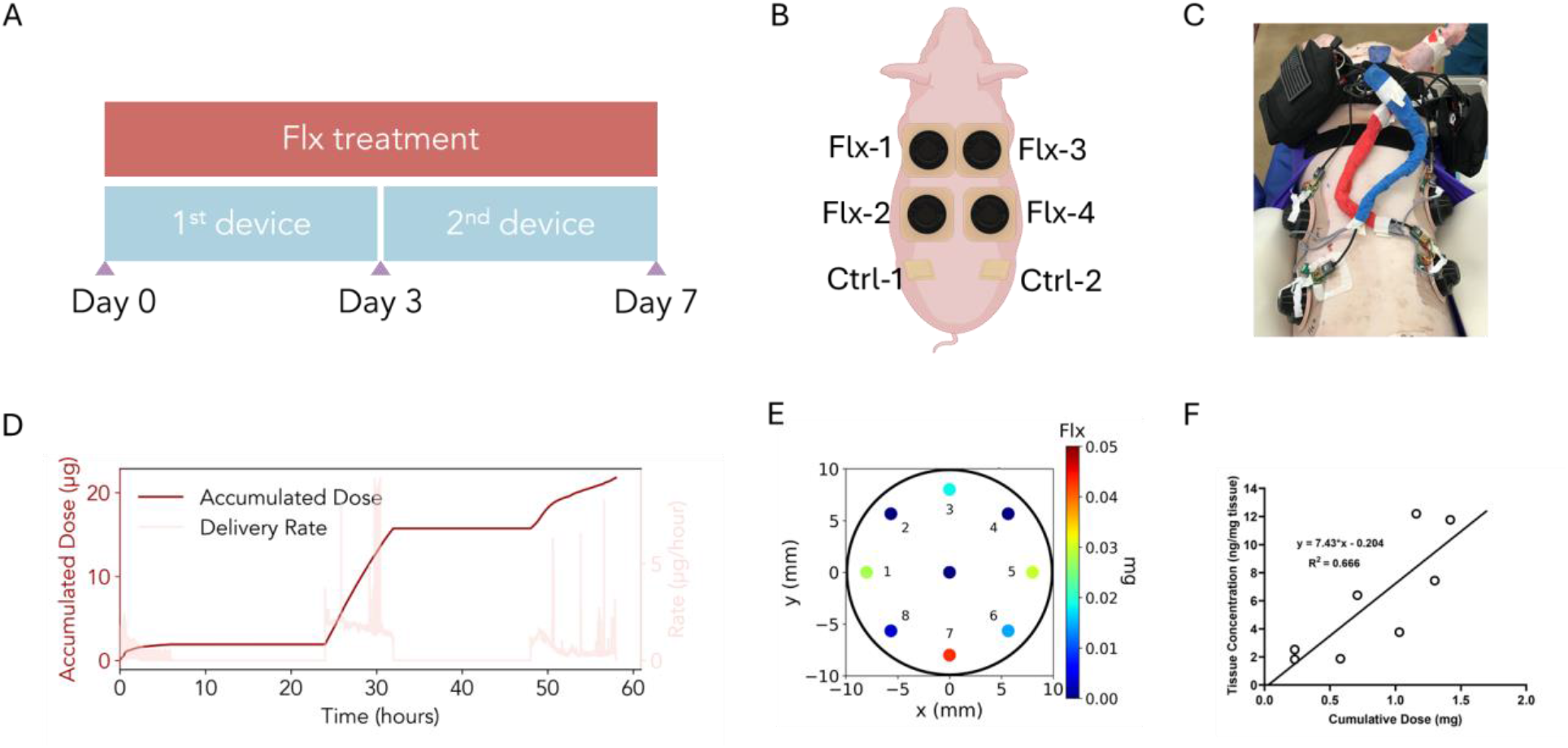
In vivo performance of the bioelectronic device. (A) Experimental schedule for 3-day and 7-day experiments evaluating short-term and extended performance. (B) Experimental setup for mounting the bioelectronic device on porcine models. (C) Image showing the bioelectronic device securely mounted on the animal model. (D) Measurements of fluoxetine delivery rate and dose during the experiment. (E) Dose map of the wound showing the spatial distribution of therapeutic delivery. (F) measured local fluoxetine concentrations from wound tissue VS cumulative dose of fluoxetine from the bioelectronic delivery.

The bioelectronic device was securely mounted onto full-thickness wounds on porcine models, as shown in Figure 4B and 4C, demonstrating seamless application and integration. The programmable fluoxetine delivery system ensured sustained therapeutic exposure throughout the experimental period. Each delivery protocol was designed to maintain consistent dosing tailored to optimize healing conditions. Simultaneously, the integrated imaging module captured high-resolution images of wound progression (Figure 4F), enabling experts to evaluate wound conditions through detailed visual data.

The results demonstrated significant improvements in wound healing outcomes, supported by dose mapping of the wound (Figure 4E) and consistent device performance in delivering fluoxetine (Figure 4D). Inflammation was reduced, as evidenced by decreased redness and swelling at the wound edges, while the center showed enhanced tissue granulation. The combination of precise therapeutic delivery and continuous imaging provided a robust platform for evaluating and optimizing treatment efficacy. This in vivo validation underscores the potential of the bioelectronic device as a versatile tool for advanced wound care in clinical and experimental settings.

The results of this study also demonstrate the efficacy of the developed bioelectronic device integrated with an onboard camera for monitoring and enhancing wound healing. The multifaceted impact of the treatment, highlighting accelerated tissue repair, modulation of inflammatory responses, and improved neuronal regeneration is illustrated in Fig. 5. Real-time imaging enabled continuous assessment of wound conditions and provided high-resolution data for evaluating the healing process (Fig. 5A).

**Figure 5.**
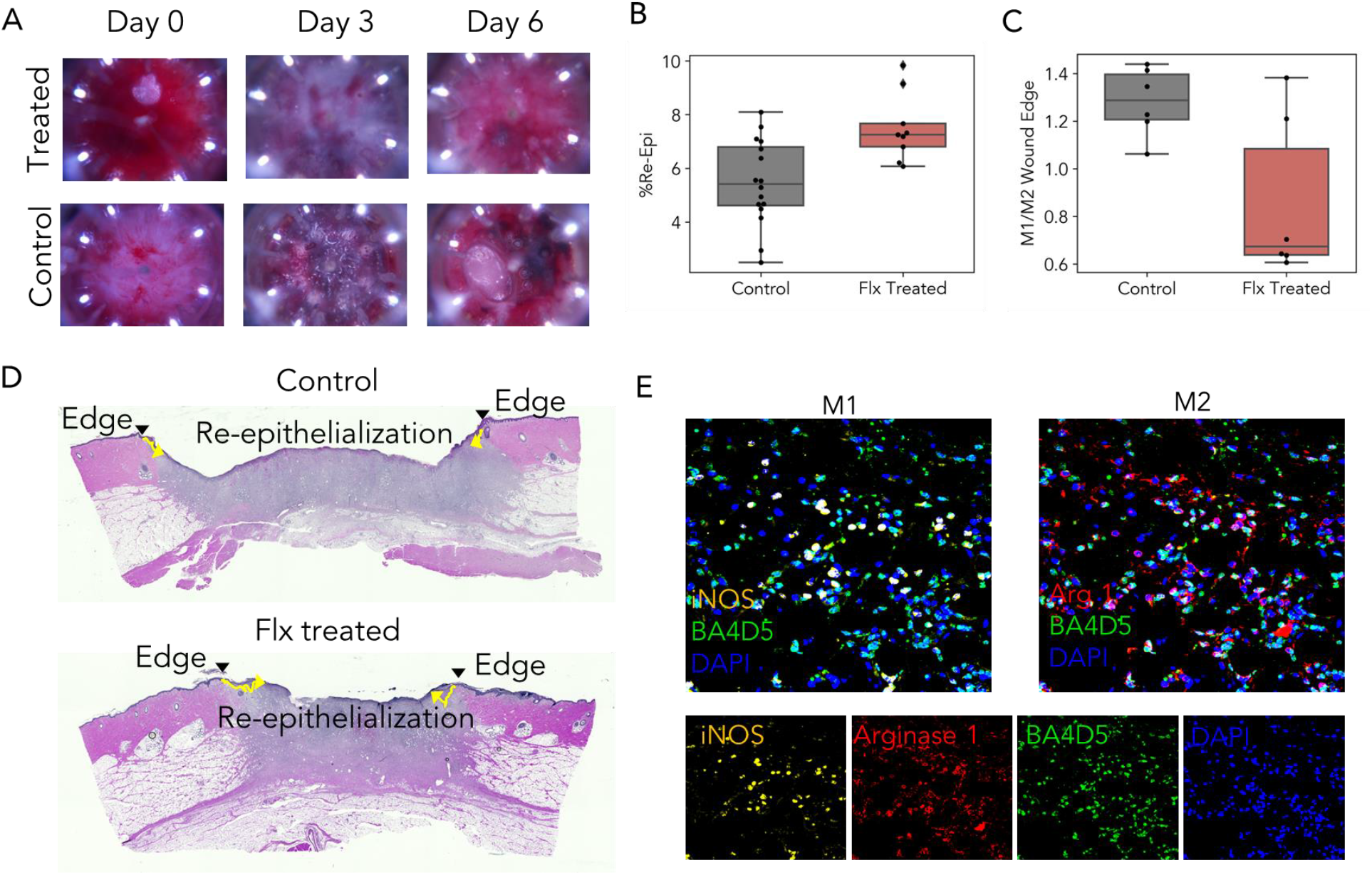
In vivo analysis of wound healing outcomes. *(A) High-resolution images of wounds captured by the bioelectronic device’s onboard camera, providing real-time monitoring of wound morphology. (B) Histological* results *of wound tissues using H&E staining, showing* 37% increase in *re-epithelialization in treated wounds compared to controls on days 3(n*_*control*_ *= 16, n*_*treated*_ *= 9, p < 0*.*01). Quantitative analysis confirms significantly accelerated healing in the treated group. (C) M1/M2 macrophage ratio demonstrating a* 33% *reduced* M1/M2 *ratio in treated wounds(n*_*control*_ *= 6, n*_*treated*_ *= 6, p < 0*.*03)*, i*ndicative of decreased inflammation and a shift to the reparative phase of healing. (*E) Staining of M1 (iNOS), M2 (Arginase 1), and the pan-macrophage marker BA4D5 showed a 33% reduction in the M1/M2 ratio in treated wounds compared to control wounds (n_control_ = 6, n_treated_ = 6, p < 0.03).

Histological analysis using Hematoxylin and Eosin (H&E) staining revealed a marked improvement in re-epithelialization in treated wounds compared to controls. By day 3, the treated wounds exhibited 37% higher re-epithelialization(n_control_ = 16, n_treated_ = 9, p < 0.01), highlighting the accelerated tissue repair achieved through the treatment (Fig. 5B, 5D).

To understand the impact of the treatment on the inflammatory response, we analyzed the M1/M2 macrophage ratio, a critical indicator of the wound healing stage. M1 macrophages drive pro-inflammatory responses, whereas M2 macrophages are associated with tissue repair and regeneration. Staining with iNOS(for M1), arginase 1 (for M2), and BA4D5 pan-macrophage marker showed a 33% lower M1/M2 ratio in treated wounds(n_control_ = 6, n_treated_ = 6, p < 0.03), suggesting a faster transition to the reparative phase of healing (Fig. 5C, 5E). Neuronal regeneration, a key aspect of functional recovery, was assessed using PGP 9.5 staining. Treated wounds showed a trend of enhanced nerve growth by day 3 compared to controls, indicating the treatment’s role in supporting both structural and functional restoration of the injured tissue (Fig. S 1).

These findings collectively highlight the capability of the bioelectronic device to not only monitor wound conditions but also facilitate key biological processes involved in tissue repair, inflammation resolution, and neuronal regeneration.

## Discussion

This study demonstrates the potential of bioelectronic systems for advanced wound care, focusing on the controlled and programmable delivery of fluoxetine through an ion pump. The device was successfully tested in large animal models, such as pigs, which closely mimic human wound healing dynamics. This achievement marks an important step in validating the system’s capability for clinical translation.

The ion pump-based platform enabled sustained and targeted delivery of fluoxetine directly to the wound site. This precise control ensured consistent therapeutic exposure, resulting in significant improvements in healing outcomes. Histological analysis revealed 37% higher re-epithelialization observed by day 3 compared to standard-of-care controls. Additionally, the device effectively modulated the inflammatory response, as evidenced by a 33% reduction in the M1/M2 macrophage ratio, indicating a transition from a pro-inflammatory state to a reparative one. Beyond inflammation reduction and tissue repair, the treatment facilitated neuronal regeneration, a critical aspect of functional recovery. PGP 9.5 staining showed a trend in increased nerve growth as early as day 3, showing the broader therapeutic benefits of controlled fluoxetine delivery.

Compared to conventional wound care methods, this system offers significant advantages as the remote-controlled ion pump delivers fluoxetine precisely where and when needed, ensuring optimal therapeutic effects. The programmable and targeted nature of the device not only accelerates tissue regeneration but also reduces inflammation and promotes functional recovery. By successfully testing this platform in a porcine model, this study highlights its feasibility and efficacy for wound healing in large mammal models that are closer to humans.

## Acknowledgments

AI assisted technologies (ChatGPT) were used to revise the draft manuscript for current grammar and improved readability of the content written by the authors.

## Materials and Methods

### Fabrication of the PDMS Actuator

The PDMS actuator was fabricated by molding 3 key components: the reservoir (76.1mm^3), the notch, and the channel (39.26mm^3). Molds for these components were created using 3D printing (Formlabs 3B) and CNC machining (Carbide 3D Nomad3). A 50-micron layer thickness was used during 3D printing to ensure precision, while CNC-milled acrylic was incorporated to provide optical clarity for imaging purposes.

The PDMS precursor was prepared by mixing the base and curing agent (Sylgard 184) in a 10:1 ratio. The mixture was degassed to remove air bubbles and poured into the prepared molds. After leveling the material for uniformity, the molds were cured at 60°C for 48 hrs. Once cured, the components were removed from the molds and cleaned with isopropanol, rinsed with distilled water, and air-dried to ensure cleanliness and readiness for assembly.

### Integrating Electronics and Coating

Silver wires (0.37mm Diameter Ag wire, Advent Research Materials) were inserted into the PDMS reservoirs to function as the working electrodes and were connected to the controller module. The connections were sealed with PDMS to prevent leakage. The device assembly was performed using clamps, and the components were bonded through oxygen plasma treatment (50W for 20 @ 0.05 Torr) to form the PDMS body with microfluidic channel and reservoirs for ion pump. After bonding, we apply either a pressure or vacuum created by a syringe to verify the seal, ensure device integrity.

A parylene coating (1um) was applied outside the PDMS piece to protect the device and enhance its durability. During the coating process, adhesive tape was used to shield the optical windows (20mm, preserving their functionality for imaging purposes (Arducam 64MP).

Afterward, the pins were connected to the controller module, which is a custom-designed printed circuit board (custom designed and send to PCBWay, Inc. for manufacturing). Following this, a chlorinated silver wire was incorporated as a counter electrode, and an additional parylene coating layer (1um) was applied to further improve device robustness.

### Final Assembly and Testing

To avoid contamination, the final assembly happens in a sterile environment in biosafety hood. Glass capillaries (1mm OD 0.8mm ID, CTechGlass) were etched with NaOH and treated with silane A174 to enhance surface compatibility. These capillaries were then filled with hydrogel precursor solutions, formulated as 1 M of 2-acrylamido-2-methyl-1-propanesulfonic acid (AMPSA), 0.4 M of polyethylene glycol diacrylate, and 0.05 M of photoinitiator (I2959), and cured under UV light (365 nm UV light at an intensity of 7.1 mW/cm^2^), following previously established protocols[19]. The cured capillaries were cut into 5-mm segments and loaded with specific solutions (10mM FlxHCl).

The prepared capillaries were inserted into the designated channels of the PDMS structure, and the assembly was sealed using a custom 3D-printed cap (19.6 mm ID) to ensure a secure fit. The fully assembled device underwent a functionality test that operates ex vivo in a clean saline solution while monitoring the current through the wifi. The device and enclosure was then glued to a skin barrier (Hollister New Image™ Flat CeraPlus™ Skin Barrier – Tape,11204) which come with adhesive that can mount the device to the wound. The devices were sealed in autoclaved stainless steel bins to allow transportation to surgery room under sterile enviroment.

### Ex vivo testing of the wearable bioelectronic bandage

The wearable bioelectronic bandage was tested ex vivo in PDMS wells filled with Steinberg solution to mimic the biochemical environment of tissue. The testing involved connecting the wearable bioelectronic bandage to a potentiostat (Metrohm Autolab) and controlling the voltage pattern via a computer while the ion pump outlet contacted the solution in the well. After a specific duration of actuation, voltage and current were recorded, and the test was stopped to collect the solution in the well. The total charge that went through the circuit was calculated by integrating the current over time, and the dose of fluoxetine was calculated by HPLC-MS. The efficiency of the delivery = moles of fluoxetine/moles of charge.

### Measurement of Fluoxetine with HPLC-MS

To measure fluoxetine concentrations in solution, we employed HPLC-MS (Thermo Scientific™ LTQ) with a reversed-phase column (Synergi™ 4 µm Fusion-RP 80 Å, 150 × 2 mm, 00F-4424-B0). A standard curve was first generated using samples with known fluoxetine concentrations. These standard samples were analyzed via HPLC-MS to confirm the fluoxetine peak based on retention time and mass readings, and the corresponding mass spectrometry intensities were recorded.

The peak area of the mass spectrometry intensity was plotted against the concentration, and the data were fitted to a calibration curve to determine the slope and intercept. For samples with unknown concentrations, HPLC-MS analysis was performed to record their fluoxetine intensity (Fig. 3D). The sample concentrations were then calculated using their peak areas and the calibration curve.

### Preparing and Wounding Pigs

Female Yorkshire-Landrace-Duroc pigs (30–55 kg) were acclimated for one week prior to the procedure. During this period, the animals were trained for handling, saliva collection, and wearing a harness to minimize stress during experimentation. On the day of surgery, the pigs were anesthetized, and their vital signs were continuously monitored. Blood samples were collected, and the surgical site was thoroughly cleaned and sterilized.

Six partial-thickness wounds, each 20 mm in diameter, were created on the dorsal surface of each pig. These wounds were either treated with standard care or equipped with Wi-Fi-connected bioelectronic devices. For device-treated wounds, the bioelectronic devices were securely mounted and powered by a portable system carried by the animal. Post-operative care included the administration of analgesics to ensure pain management and facilitate recovery.

### Monitoring and Tissue Collection

Daily inspections were conducted to monitor the condition of the wounds and the functionality of the bioelectronic devices. Dressings and devices were replaced every 3–4 days to maintain sterility and ensure optimal device performance throughout the experimental period.

At the conclusion of the study, final blood samples were collected from each pig for biochemical analysis. The animals were then humanely euthanized in accordance with ethical guidelines. Tissue samples from the wound sites were harvested and preserved for detailed histological and molecular analyses.

### H&E staining

5 um thick Tissue sections were prepared and stained with Hematoxylin and Eosin (H&E) to evaluate key indicators of wound healing, including re-epithelialization, collagen deposition, neuronal growth, and macrophage activity. High-resolution imaging (BZ-9000 inverted microscope, Keyence, Osaka, Japan) was performed on the stained sections, and BZ-II Analyzer (Keyence, Osaka, Japan) was used to quantify healing progress by measuring the extent of new skin growth over the wound areas.

### M1M2 and nerve staining

Formalin-fixed, paraffin-embedded (FFPE) skin sections from day 23 were used for immunohistochemistry (IHC) staining to visualize macrophages, neurons, and angiogenesis. Macrophages were labeled with antibodies against iNOS (M1; ThermoFisher, Catalog # PA1-036), arginase 1 (M2; ThermoFisher, Catalog # PA5-18392), and a pan-macrophage marker (BA4D5; Bio-Rad, Catalog # MCA2317GA). Multicolor labeling was achieved using donkey anti-goat Alexa Fluor 568 (Invitrogen, Catalog # A-11057), donkey anti-rabbit Alexa Fluor 647 (Invitrogen, Catalog # A-31573), and donkey anti-mouse Alexa Fluor 488 (Invitrogen, Catalog # R-37114). Neurons were identified in consecutive skin sections of 30 µm thickness using the PGP9.5 antibody (ThermoFisher, Catalog # 480012) and donkey anti-mouse Alexa Fluor™ 488 (Invitrogen, Catalog # R37114) as the secondary antibody. All slides were mounted with VECTASHIELD® Antifade Mounting Media with DAPI (Vector Laboratories, Catalog # H-1200-10) for nuclei visualization. Imaging was performed with a Zeiss LSM900 Confocal Laser Scanning Microscope at a resolution of 3 pixels per µm. For each wound sample, three stitched images of the wounded region near the healed epidermis were collected using a 40× oil immersion lens to minimize tissue variation. Quantification of macrophages, neurons, and blood vessels was performed by visualizing the staining in the confocal images, and analysis was carried out accordingly.

### Fluoxetine Concentration Measurement from Wound Tissue

Pig wound tissues (20–30 mg) were cryo-pulverized in liquid nitrogen and resuspended in 0.250 mL methanol containing 0.5% (v/v) formic acid. The samples were sonicated for 15 minutes in a sonicating bath to ensure thorough homogenization. The homogenates were then centrifuged to separate the supernatant.

The supernatant was diluted to 0.500 mL by adding 0.250 mL water. Finally, 10 µL of the prepared extract was injected into the analytical system for fluoxetine concentration analysis.

## Supplementary Information (SI)

### Calculation of fluoxetine dose

Fluoxetine was delivered using a bioelectronic ion pump, as explained in previous studies[20]. In this system, fluoxetine becomes protonated in slightly acidic conditions (pH ∼6), turning into a positively charged ion (Flx^+^). These Fl^x+^ ions then move from the working electrode to the ground electrode due to an electric field, a process called electrophoresis.

Since Flx^+^ is the main ion in the source reservoir, the amount of fluoxetine delivered is related to the total electrical charge (the flow of electrons) passing through the circuit. To measure how efficiently the device delivers fluoxetine, we used HPLC to create a standard calibration curve. By tracking the current over time, we calculated the total charge, which helped us determine the delivery efficiency of the device.

### Detection of fluoxetine in pig plasma

To check if the fluoxetine delivery would cause any systematic result, we used HPLC to detect fluoxetine concentration in plasma of the animal. Result shows fluoxetine was not detectable in pig plasma (n=3 pigs) following topical fluoxetine wound treatment using the experimental device. Animals received topical fluoxetine on four wounds each day over the course of seven days, and plasma was collected one day after the final dose of fluoxetine was administered.

**Fig. S 1.**
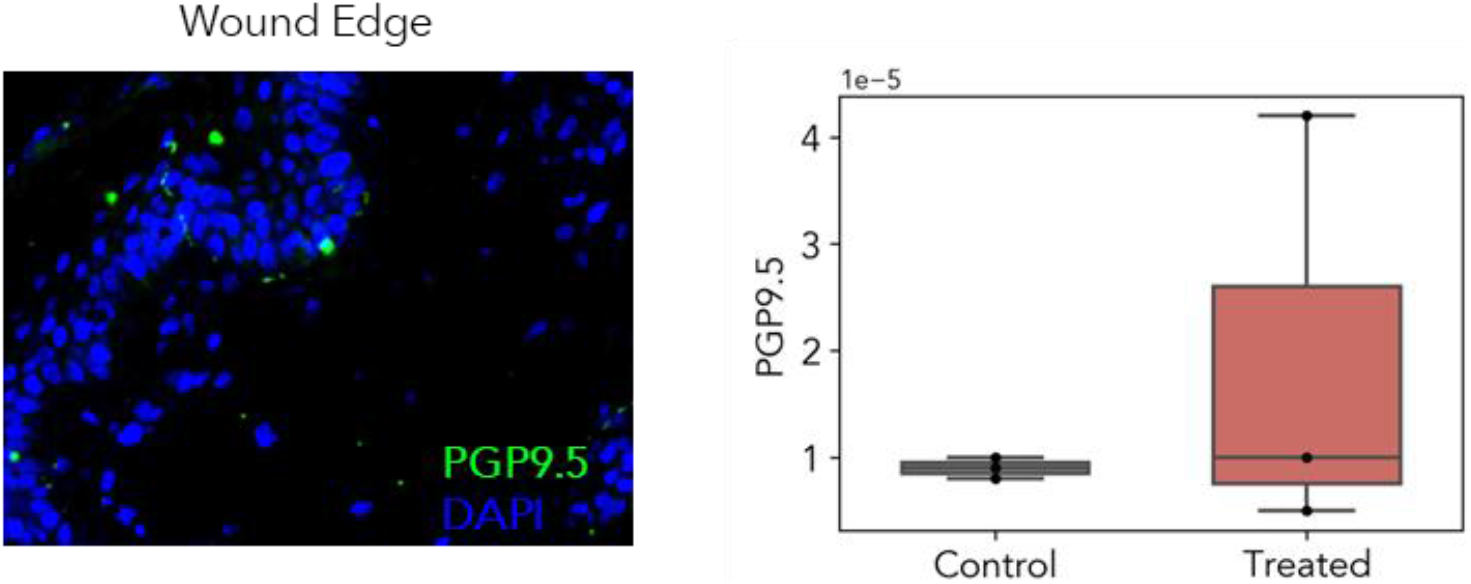
PGP 9.5 staining reveals 111% increase(n_control_ = 3, n_treated_ = 3, p = 0.48) in neuronal regeneration in treated wounds by day 3, show a trend in promoting nerve growth and functional restoration.

